# 3D Direttissima: A V1 Heresy

**DOI:** 10.1101/2022.08.10.503492

**Authors:** David W Arathorn

## Abstract

The heretical conjecture offered here is that a 3-space scene reconstruction of sequences of visual fixations is constructed directly in V1 in a head-centered frame. “Direttissima” is a climbing term designating the shortest, most direct route to the objective of a climb. The “heresy” here is the claim that the construction of the first 3D representation of the scene before one’s eyes takes place in V1 rather than further up the visual pathways. The inputs to the neural process proposed for such construction are the V1 retinotopic projections from each eye and the vergence angle. Absolute disparities are not encoded as a separate data set in a distinct set of neurons, but rather are emergent properties of the neurons that implement the construction of the 3D representation of one or a sequence of fixations on elements of the scene. In this conjecture, the conscious “picture” we take as vision is a projection of the 3D construction, not a view of the retinal image. One of the critical psychophysical clues to the nature of such a mechanism is the well known visual size constancy phenomenon. Another property of the proposed mechanism is that it is plausibly fast enough to enable such spatially and temporally demanding tasks as a human batting a fastball or a bobcat swatting the head of a striking rattlesnake.

## Introduction

The heretical conjecture offered here is that a 3-space representation of sequences of visual fixations is constructed directly in V1 in a head centered frame. The inputs to this construction are the binocular retinotopic “images” in V1 and the pointing angles for each eye. Separate encoding of absolute disparities is not an input, but rather image disparities emerge as part of the 3-spatial construction. The result of the spatial construction is a neural equivalent of a point cloud in machine vision. Because, in this conjecture, the 3D construction takes place In V1 directly from the retinotopic 2D representations in V1, the 3-space representation can arise at high speed. Along with the 3D representation of the visual scene, another output of this neural circuitry is a 2D projection of the 3 space in which separate objects exhibit the size constancy characteristics long recognized in visual psychophysics [1], which contradict the veridical size relationships that exist in the retinal image and retinotopic V1 representations.

A critical characteristic of any system which produces such a 3-space representation is that within that 3-space construction the location of observed static objects must be constant and independent of the momentary direction of gaze and vergence distance. This is a requirement in order that a sequence of saccades across a scene does not produce conflicts or instabilities within the 3-space construction. In this discussion we will assume constant head orientation during the period of construction, though the more general case is readily accommodated.

The mechanics of such a system are presented here in a highly idealized graphical, non-mathematical, form. The idealization sets forth the basic functional requirements for a neuronal implementation, but no specific implementation is suggested. The principles of operation are the intended target of the presentation here.

The guiding constraints, therefore, are:

1. 3-space construction must be consistent independent of multiple retinal frames of reference used as input.
2. Vergence is the only depth cue used. No object knowledge or top down influence is used in this stage.
3. Infinite object distance must be represented within a finite (neuronal) 3-space.
4. An output of the system must be a 2D scene representation which exhibits size constancy characteristics consistent with established psychophysics.
5. Must require no higher level encodings of retinal inputs than are known to exist in V1.
6. Processing must be parallel, feedforward, and non-iterative to be consistent with performance known to be required for such tasks as fastball location.

### The Conjecture

The diagrammatic formalism used to present the mechanism needs some explaining. The wide rectangle represents the neuronal “zone” while the area above it represents the optical zone. The two small circles at the interface between these zones represent the eyes, and the heavy double headed arrows through the eye circle axes represents the retinotopic projection plane, i.e. in this abstraction, the image plane of each eye. The pointing direction of each eye is the perpendicular to the projection plane in the optical zone.

In the neuronal zone the rectangle represents two adjacent planes, one for each eye, of a 3D volume of neurons in which the 3D reconstruction of the scene will reside. The two planes are represented side by side in the diagram, though the more likely physical arrangement is stacked as alternating layers or slices since that architecture minimizes the length of the interconnections between the adjacent layers for left and right eye.

The heavy horizontal arrows in the optical zone represent the front surface of physical objects. For the eyes, pinhole optics are assumed, except that in the neuronal zone the rays that would be optical are sequences of neurons spaced along the continuation of each optical ray. The adjacent planes are continuations of the same optical zone plane for each eye.

The neuronal volume becomes the residence of what amounts to a “point cloud” representation (as in machine vision usage) of the scene before the eyes. For those readers not familiar with the term, a point cloud is constructed from a specialized 2D image which records both the illumination and distance from camera to imaged surface for each pixel in the image. Since the image formation parameters for a depth camera are fixed, it is simple to invert the projection from the 3D world to the image plane of the camera to create a 3D representation of the visible surfaces of the objects in the world which formed the image. In a machine vision point cloud, the inverse projection renders each object’s surface at a fixed scale relative to its size in the physical world, unlike the 2D image, in which the size of each imaged object in inverse proportion to its distance from the camera. (This property has important consequences in the proposed biological visual system.) Each point in the machine vision point cloud is a tuple of illumination and x, y and z spatial coordinates. This data set is kept as a list for compactness since there are only as many defined points in the cloud as there are pixels in the 2D image. However, the point cloud could just as well locate each illumination datum from the 2D image at the appropriate <x, y, z>th element of a 3D array, leaving the vast majority of elements having zero value. The neuronal architecture proposed here corresponds to the latter explicit 3D array representation, with a neuron or several to hold the value of the illumination at the implied 3-space coordinate.

The fore-aft axis of the neuronal volume becomes a depth mapping of the distance from the eye plane to the objects being imaged. The neuronal circuitry located within the neuronal zone performs the inverse projection from retinal image plane as represented in V1. Since, unlike the data from a depth camera, there is no explicit datum for the depth of each “pixel” the depth computing neural circuitry depends on the parameters of vergence and the spatial relationship of corresponding image features in the 2D projections of each eye, usually referred to as disparity.

The diagonal lines into the neuronal region emanating from each eye represent the the inverse projections for the current eye direction from each “pixel” of the 2D image plane. Since these inverse projections are indeterminate in depth each intercepts all neurons in the region which lie along that ray. Each can be idealized as a single long axon synapsing on the dendrites of the 3D array neurons which lie on, or very near, its path. However, which of those neurons on its path it activates depends on additional circuitry from the corresponding region of the 3D array for the other eye. That circuitry tests for feature matches over local regions with a geometrical offset between the eyes corresponding to horizontal disparities in the 2D images. There is an additional input to that circuitry, which is an encoding of the vergence angle. Vergence and disparity, as usual, determine depth, so the circuitry associated with each neuron in the “point cloud” array, determines from those parameters which neuron along the radiating axon from the 2D image plane becomes activated as the representative point in the 3D cloud.

Only the result of that depth-determining process is addressed here. The specific circuitry which might implement it is left to another presentation, or to the reader’s ingenuity.

There is another complication arising from the finite size of the 3D array. In a machine vision point cloud, objects lying at great distances from the camera are processed as if lying at infinity. However, whether the depth camera determines depth from time of flight or stereopsis on a projected IR pattern, the camera has a limited depth range that it can encode. Pixels in the camera’s illumination image from objects at distances exceeding the depth range will show up normally in the RGB image, but in most implementations will have no depth value associated with them, so in fact lie outside the limited point cloud maximum dimensions. Since depth in this proposed architecture is solely determined by disparity and vergence there is a limit to meaningful depth computations. It is possible that the biological implementation also “clips” out-of-range depth. But it is also possible that it represents depth on a non-linear scale which places objects at distances beyond meaningful depth computation inside the limit of the neuronal 3-space array. There are several psychophysical phenomena which suggest the latter. First is our tendency to increasingly underestimate distances to objects as they are located further from us. Second is the perceptual geometry of the size constancy phenomenon. For those reader not familiar with this psychophysical characteristic, it has long observed that viewed objects do not decrease in perceived size as a function of distance as they do in a photographic image, or in the optical retinal projection. Instead, objects appear at a size they would appear if much closer than they actually are. And yet we do not misjudge their physical distance from by anywhere near as much as their perceived size would suggest. This means that their actual distance is fairly accurately represented, particularly within reaching distances, and yet their perceived size is not consistent with what optical law predicts for that distance.

The above is the reason for the curved contour lines in the diagrams. In machine vision point clouds, the dimension of the representation of the surface of an object is proportional to the true size of the object, as mentioned above. To illustrate, assume a point cloud is constructed from an image of two similarly sized apples, one a distance *d* from the camera and the other at distance 2*d*. In the 2D image the further apple appears to have half the diameter of the nearer ball. But if one were to make a 2D parallel projection along the z (depth) axis of the point cloud constructed from the original 2D camera image, both apples would appear to have the same diameter, though the further will be more sparsely pixilated since the further ball contains ¼ of the pixels compared to the nearer in the 2D camera image. The parallel machine vision point cloud projection would exhibit absolute size constancy, independent of distance. Our biological visual system exhibits relative size constancy, in which the more distant apple appears reduced in diameter but by far less than it ought. This phenomenon can be simulated by computing a converging projection from the point cloud along the z axis. With a converging projection, objects with greater distance along the z axis from the projection plane appear somewhat smaller. In Figure 2(a) two nearly same-size apples were imaged by an Azure Kinect depth camera, one at approximately 50cm and the other at approximately 100cm. The original photographic image (filtered for apple red) shows the expected 2 to 1 diameter relationship. A slightly converging projection of the point cloud, Figure2(b) shows relative size constancy as is observed (approximately) by a human from the camera position.

The converging contours in the diagram implement this relative size constancy. Each contour represents where the feature matching circuitry actually locates the activated neuron for the axon carrying the retinal projection “pixel.” The contours specify that the distance representations (along the z axis) are always correct, but the perpendicular dimensions (x and unillustrated y axes) decrease with increased distance, but a rate consistent with the human psychophysics of size constancy. Independent of the foregoing, z axis distance encoding is non-linear, such that nearby distance deltas are given more z distance in the 3D array than far distance deltas. This encoding allows objects at optical infinity not to be clipped by lack of depth encoding range in the array. And unlike the machine vision point cloud, in which objects at great distances are not represented, in this representation the brain equipped with such a mechanism can readily assign “far away” to objects encoded near or on the back boundary of the 3D neuronal array.

The contours also specify how by how much the each point in the reconstruction for each eye has to be shifted to reconcile the disparity differences between the two eyes and thereby construct an accurate (but scaled) 3D reconstruction which preserves the original alignment of the objects being viewed. In Figure 1 this depicted by the centrally shifted arrows (now rendered as heavy dotted arrows) from the 3D reconstruction for each eye. Notice this preserves the relative size constancy present for each eye, but corrects the disparity shift so that the butt ends of the arrow are aligned on the central axis, as they are in the real scene. The box at the bottom of the diagram contains what would be the projection of this disparity reconciled reconstruction. It is also shown in Figures 4 and 5. The disparity reconciled reconstruction is an alternative source of conscious vision (as will be discussed later) because this is where partially occluded views of an object in each eye would be merged or fused into a whole to produce that known visual competence [2]. The disparity reconciled reconstruction provides an excellent source for remapping to a dimensionally linear, size-correct 3D representation useful for reach and grasp. However, the common experience of “double” vision of objects further or nearer than the point of vergence persisting along with relative size constancy argues for the separate left and right eye reconstructions as a source of conscious visual imagery. According to [2] whether binocular rivalry or merging predominates depends on object surface characteristics. This discussion simply offers that both percepts can emerge from the conjectured mechanism, but not how conflicts would be resolved.

**Figure 1.**
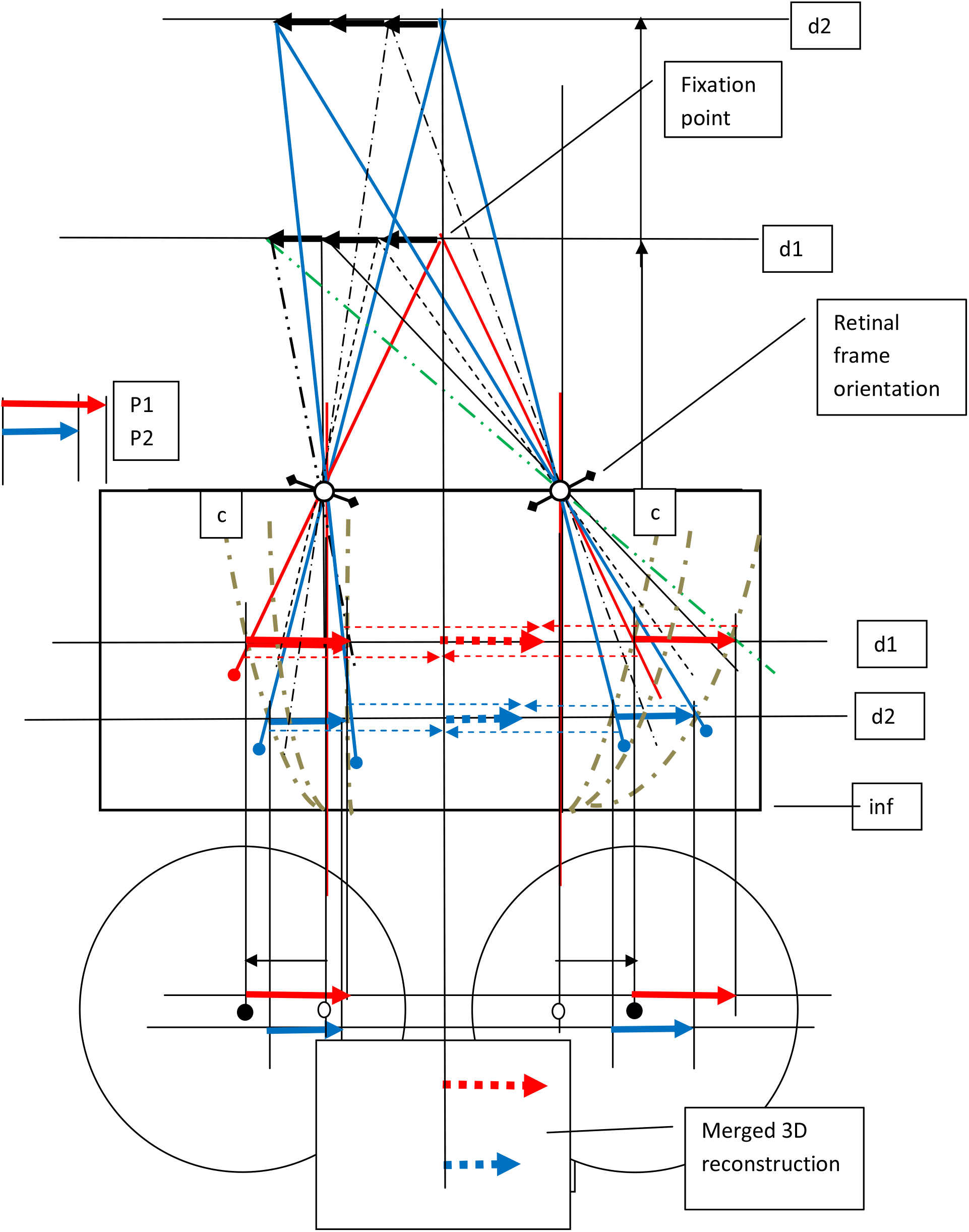

**Figure 2:**
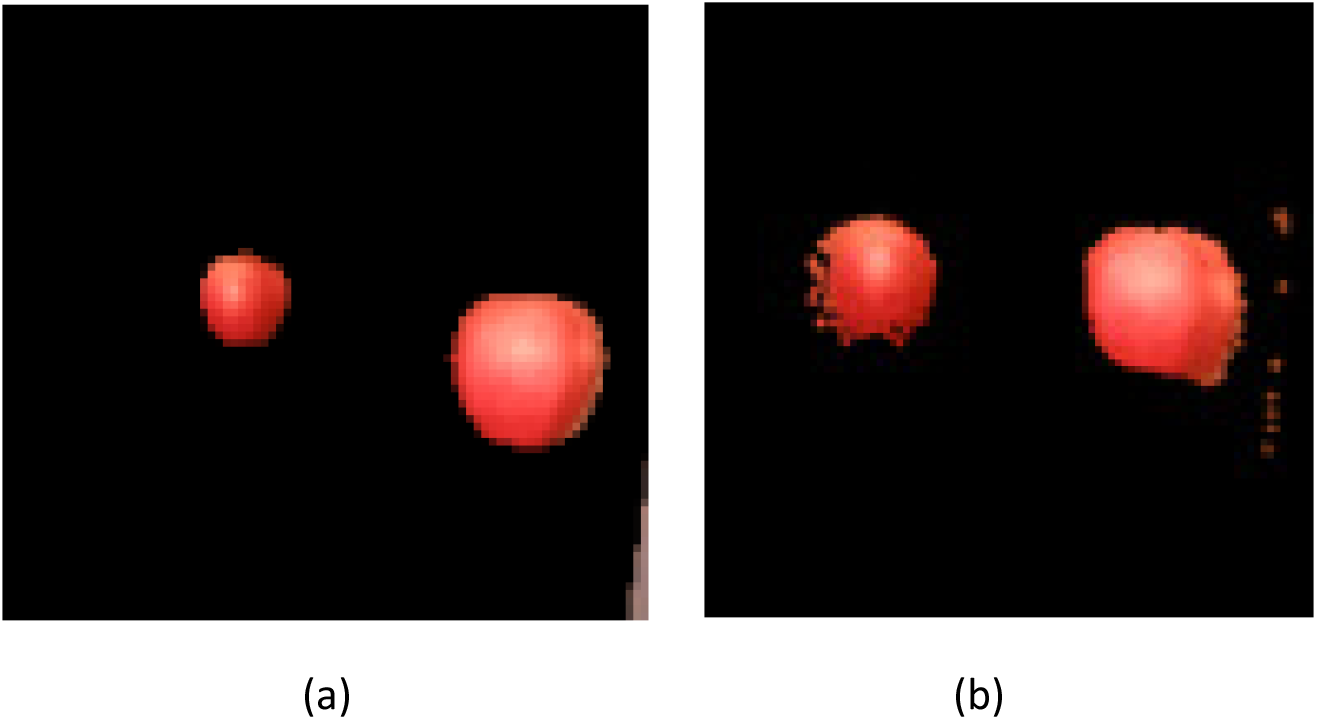
Simulation of Size Constancy Psychophysics. (a) Photographic image, diameter inverse to distance, (b) Converging projection of point cloud simulates relative size constancy as observed by humans.

It should be kept in mind that the diagrams greatly exaggerate the interocular distance relative to normal physical distances of observed objects. In other words, the diagram illustrating the nearer fixation would, if to scale, imply an extreme cross-eyed vergence to some object lying not far beyond the tip of one’s nose. This exaggeration is necessary to make the diagrams readable. But one has to visualize the horizontal dimension to be a fraction of what is shown to be in correct proportion to the depicted object distances. As a consequence the feature match contours would in reality be far more elongated and less curved.

As illustrated in Figure 1, for centered vergence at distance d1 the *c* contour for center “rendering” of the fixation point in the 3D array matches points in the image from the other eye having zero disparity. For a more distant centered object point at d2, the same contour locates the inverse projection at the d2 representation along the z axis of the 3D array, but with somewhat reduced x (and implicitly y) displacement. That locus on the contour is associated with disparity of 2*k* between the left and right eye retinal images, as illustrated in Figure 3.

**Figure 3:**
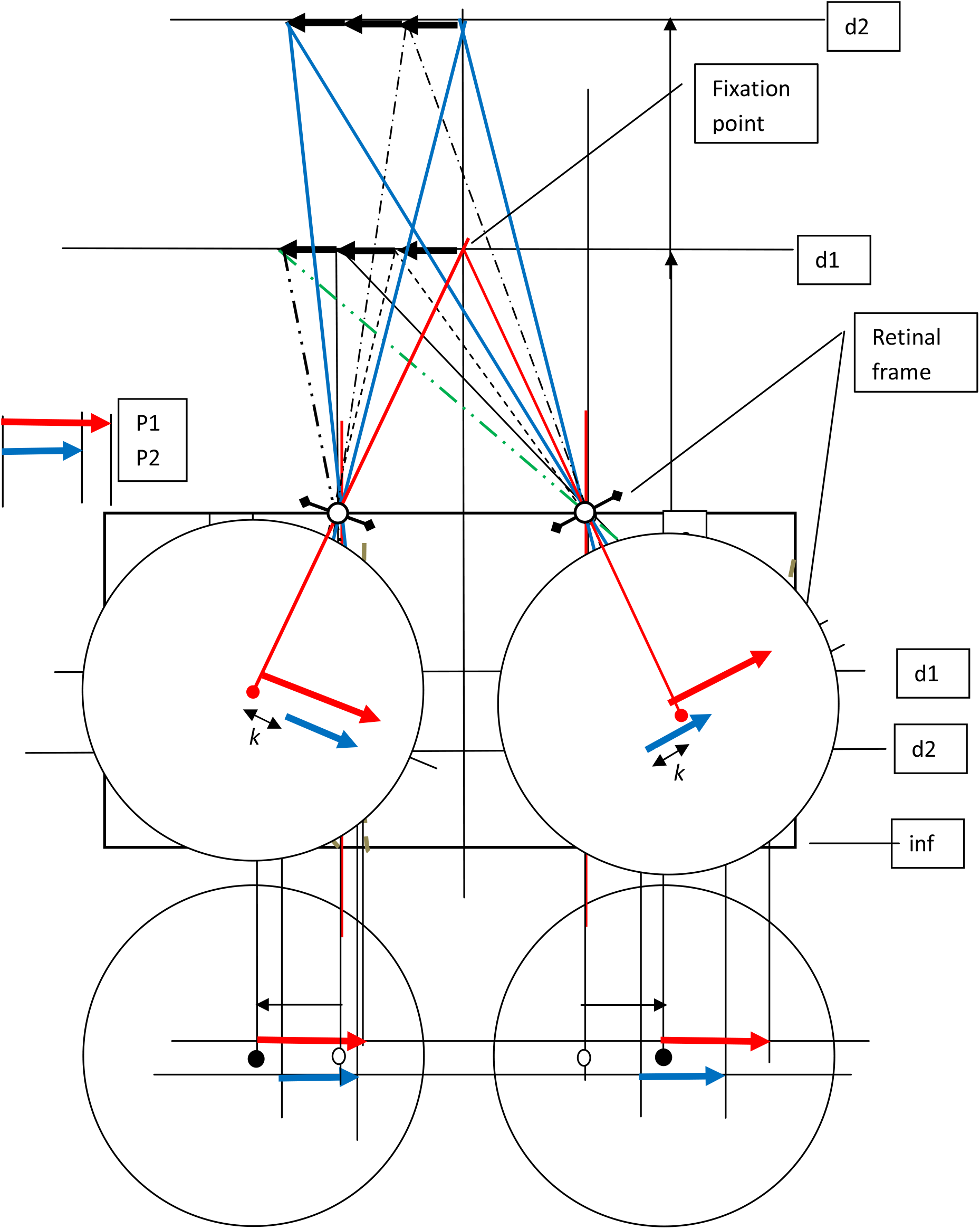
Retinal Projection versus Point Cloud Projection, near fixation.

**Figure 4:**
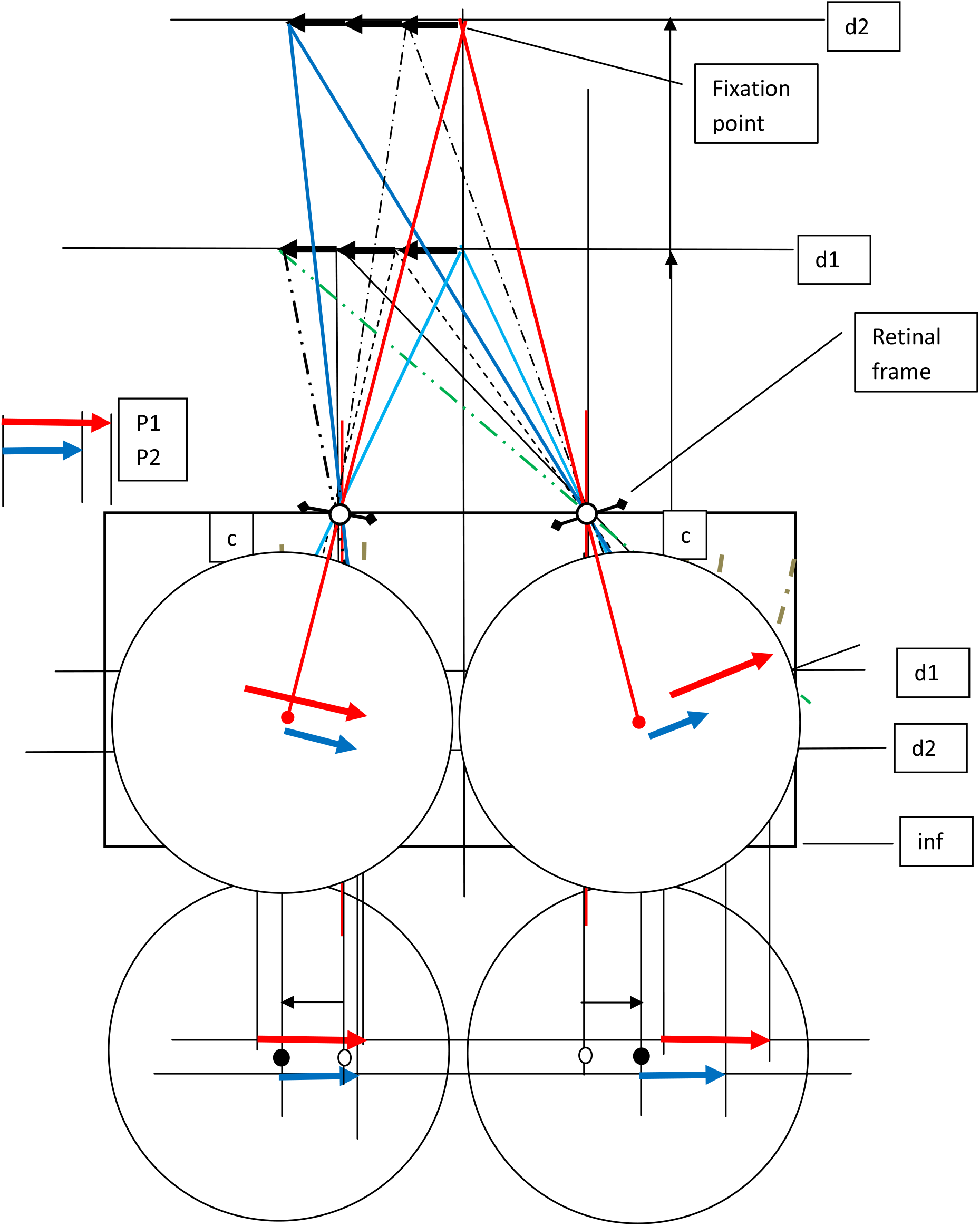
Retinal Projection versus Point Cloud Projection, far fixation.

**Figure 5.**
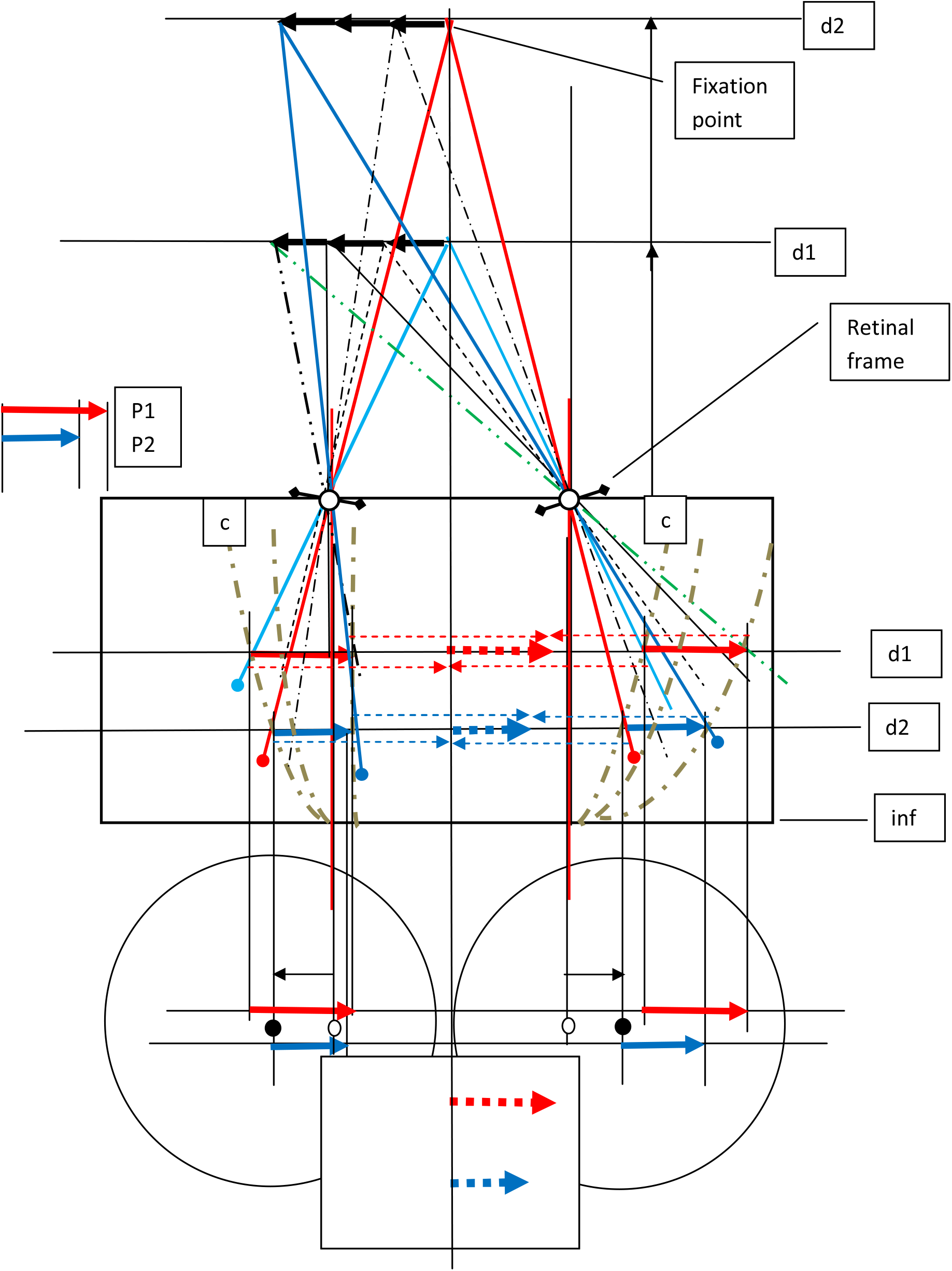

As also illustrated in Figure 1, object points off the vergence axis are rendered along contours further from the center *c* contour. For off center points on objects at the vergence distance, zero disparity associated with lateral contours at z distance d1, while for objects at distance d2 the lateral contours match retinal image disparities of 2*k*.

The fact that the center *c* contour is curved implies that object points lying along the axis through centered vergence do not lie along a straight line in the 3D array. As mentioned earlier, this is corrected in the shifted and merged reconstruction shown between the two separate eye reconstruction 3D arrays. The merged reconstruction is “veridical” with respect to central alignment, while it retains the known relative size constancy characteristics.

### What do we see?

This is an appropriate point to raise the question: what are we seeing in our “mind’s eye?” Size constancy argues convincingly that we do not “see” the retinal image in which an object’s size is inversely proportional to its distance. In the conjecture presented here, the source of the imagery that produces the observe size constancy characteristics is a 2D projection of the 3D array. Figures 3 and 4 illustrate the dichotomy of the object sizes in the retinal projection versus a projection of the “point cloud.” From a computational system perspective, making the 3D array the source of the image we see also makes it trivial to associate a distance (and general position in the 3D world, at least approximately) with anything that gets our attention in the visual field, and even objects that don’t get our attention. And, as will soon be discussed, this will be true for 3D “point clouds” constructed from multiple fixations, as typical of our pattern of visual scene acquisition. The question “what do we see” also must address the dichotomy between fused and rivalrous projections, as would emerge from the merged 3D reconstruction and the separate 3D reconstructions for each eye, as discussed earlier.

### Multiple fixations

Since we most often assemble our view of even a single object from a rapid sequence of saccades with different fixation directions, often at different distances, assembling a coherent version requires transformations from the sequence of retinal frames of reference to a single frame of reference. This too argues for that frame of reference to be a 3D frame, since the scaling from retinal images formed at different distances and therefore different scales, must be reconciled. The assembly of a point cloud in the 3D array provides a natural target for this process. The condition that must be satisfied is that the same object geometry, observed with different fixations, must produce the same 3D construction. As Figures 1, 5 and 6 illustrate, that is the case for the hypothesized mechanism described here. Figure 5 illustrates a central fixation on the more distant of the two objects shown in Figure 1. Figure 6 illustrates a fixation 2/3 of the way to the left end of the same more distant object. In all cases the same representation is constructed in the same locations (with a caveat which follows) in the 3D array.

**Figure 6.**
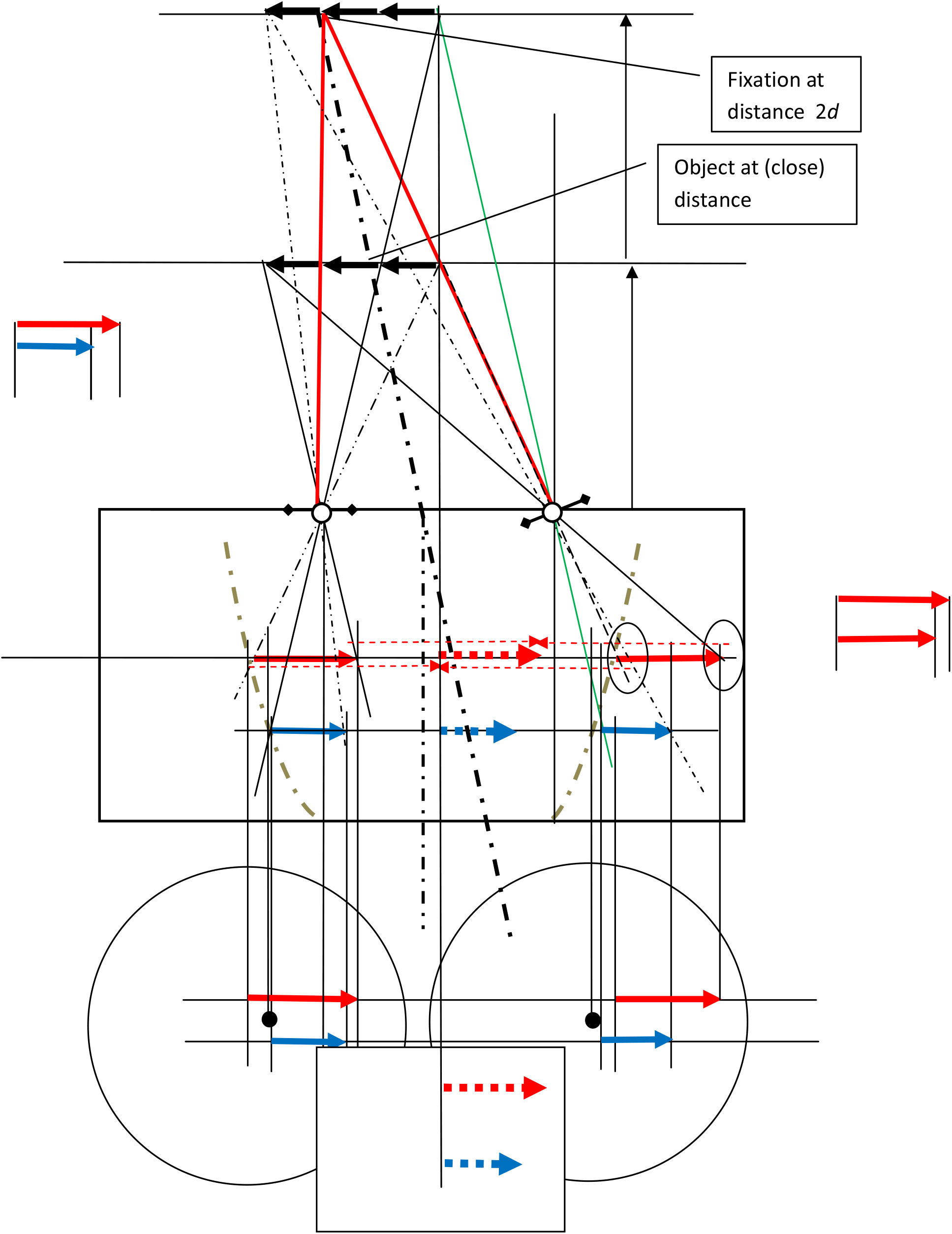

However, it should be noted in Figure 6 that there are small inconsistencies between the left and right eye 3D reconstructions, as emphasized by the small ellipses. This is due, of course, to the asymmetry and the consequent slightly different scales on each side. In a central disparity shifted reconstruction the two sides will not match exactly (at least without some small scaling operation). To preserve consistency across fixations, the inverse projection that is closest to the central angle is the one that will match the reconstruction from the central fixation.

While Figures 1, 5 and 6 illustrate the constancy of 3D construction for a variety of depth and direction of fixations for a particular object configuration, an argument for the generality of the assertion runs as follows.

Suppose the eye is an ideal pinhole camera with a spherical image formation plane concentric with the pinhole. No matter what direction an object is located in front of such an eye/camera, so long as its distance to the pinhole doesn’t change, the image of that object on the imaging sphere surface remains the same, though located at different locations on the imaging sphere. Suppose we introduce a circular mask with a shutter behind the pinhole, that restricts the diameter of the image projected onto the imaging surface. The mask is constrained to move on a spherical path concentric with the imaging surface. We keep the position of the object of interest constant and allow the mask to move around rapidly to various positions so as to sample the object and assemble over multiple exposures a picture of the whole object on the spherical imaging surface. It should be apparent that if the object being imaged is stationary, the assembled image is coherent and geometrically the same as it would have been with a single exposure without the mask.

Now imagine that the imaging surface behind the mask is fixed to the mask and rotates with it. Now the image formed at each “saccade” on the new imaging surface each has its own frame of reference, i.e. the ‘retinal” frame. This is a simplified analogy of the eye. The mask-locked images cannot be meaningfully assembled unless each one’s frame is transformed into a general frame, such as the original fixed spherical imaging surface. The latter transformation is a rotation of the mask-locked imaging surface corresponding exactly to the rotation parameters of the mask. That is a realization of what is being proposed here. In effect, the 3D array assembly mechanism treats the eye as if it were an idealized pinhole camera with a single fixed spherical imaging surface behind it and a movable mask and shutter. The images formed on this fixed surface are inversely projected into the 3D array as described earlier, incrementally over multiple saccades/fixations. This constructs a coherent 3D “point cloud” in the 3D array just as if the entire spherical imaging surface had been “exposed” at once.

### Reconciliation of Conjecture and Experiment

The long observed size constancy phenomenon is central to this conjecture. An early research paper characterizing the phenomenon quantitatively [1] appeared in 1963. Many have followed.

One of the most obvious features of the conjecture is the apparent absence of a class of neurons whose purpose is to encode absolute disparities. Neurons with such properties are well known to exist in V1. The apparent absence here is illusory. The neurons along the contours described earlier locate a point in the 3D array contingent upon the disparity between the sources of that point in the two retinal images and the vergence. If a subject is presented with binocular visual stimuli which embodies a well defined disparity, for the current vergence angle of the subject’s eyes, the neuron along the relevant contour will respond if it encounters the expected disparity. So while in this conjecture there is no separate class of neurons whose purpose is to encode absolute disparities, there is a well defined class of neurons in the proposed architecture which will show activation if presented with specific stimulus disparities. So the well documented existence of disparity-tuned neuron classes neither confirms nor discredits the proposal.

However, a study of characteristic response of V1 neurons to BOTH disparity and depth [3] reveals that there is more to the behavior of many V1 neurons than when tested for disparity response alone. To quote, “..we have observed a strong modulation of the spontaneous activity and of the visual response of most V1 cells that was highly correlated with the vergence angle.” And “Experiments were conducted to test disparity sensitivity using static random dot stereograms (RDS) at different viewing distances set at 20, 40 or 80 cm. It was found that more than 75% of recorded cells drastically changed their visual response when the distance of fixation changed, with the consequence that disparity selectivity could be present at a given distance but absent at another one.”

While the resolution of the experiments in [3] were not sufficient to reveal a selectivity as fine as implied by the hypothesis presented here, their result revealed a qualitative similarity in that many cells encountered appeared to respond to the combination of disparity, vergence and depth.

As discussed earlier, the conjectured architecture embodies are rotations (or shifts) in the V1 retinotopic projection of the retinal image to compensate for changes in eye direction. Remapping as a consequence of saccades is frequently documented, e.g [4], but the focus of the research has been primarily on remapping to the fovea of the target of a saccade, rather than the generalized rotation or shift of the retinotopic field to construct a coherent 2D scene representation. Of course, a broad remapping which brings a non-fixated image feature to the fovea will inherently accomplish the consist image shift required in the conjecture.. However, to the author’s knowledge the documented remappings do not address the broader transformation, neither confirming nor precluding it. This will have to be left as another consequence that is not documented because the experiments are not designed to observe it.

The latency or speed of 3D construction implied by this conjecture is best supported by sport-based experiment augmented by empirical inference. A study of visual tracking of pitched baseballs [5] indicate there is a saccade to the predicted location of the ball near the hitting zone which starts about 200 msec before the ball reaches the zone. This is about the same time as the bat swing is initiated. At 80mph (a slow fastball these days) the ball is travelling about 120 feet/sec, so it is only about halfway to the plate when the saccade starts. It will cross the hitting zone in about 10 msec, so under no circumstances is visual information from when the ball is actually inside the hitting zone itself useful. On the other hand, every ball player and every tennis player knows that if one does not “watch the ball” contact with the ball and the hitting instrument becomes very unreliable. Given the lateral motion of the ball in both sports, having even an accurate estimate of its location 60 feet away is not useful, because if it was, “watching the ball” would yield no benefit. So there is some useful ball-locating information that happens after the saccade to the hitting zone, and it must be acquired and acted on in well under 100 msec. The most likely explanation is that the saccade to the hitting zone puts the direction of view considerably ahead of the ball, but quickly enough that the ball is picked up in the periphery when it is nearing the hitting zone, 12 feet or less away, and it’s 3D location relative to the observer’s head is computed and a track predicted. At 120 feet/sec that means sub 100 msec to acquire location, predict path, and make the small motor adjustments required for the hitting performance consistent with “watching the ball” to the bat or racket. The visual experience of seeing the ball actually hit the bat or racket is a memory, and may be useful to tune the in-swing correction mechanism. But it is hard to reconcile the need for sub 100 msec in-loop observation and correction with 3D reconstruction anywhere but in V1. It doesn’t have to be high resolution 3D reconstruction, but it has to be fast and reasonably accurate out to 12 feet or less at least for major league batters and professional tennis players.

In the non-human animal world extreme physical behaviors which require very high speed 3-D localization are common. Videos of bobcats reliably swatting with their claws the heads of striking rattlesnakes are readily found on the web. A rattlesnake strike is typically 70msec from launch to bite. So the implication for the speed of 3-D location processing in the visual system of the bobcat is clear. It must take place early in the pathway. This example, unlike the baseball example, has the advantage that explanations by prediction requiring less stringent timing are precluded.

## Discussion

What this conjectured mechanism is and isn’t must be emphasized. It is 3-D scene reconstruction mechanism. As described it has no process for isolating single objects and extracting a 3-D representation thereof. Even an application as simple as tracking a baseball requires another step of processing, which this paper does not address. However, isolating a single fast-moving surface within the 3-D array is minimally complex compared to isolating and identifying specific objects. For example, simple subtraction of sequential 3-D snapshots isolates a moving target. Other simple means to the same end are possible. So the addition of such neural circuitry which would make possible the rapid location of a baseball or the head of a striking rattlesnake would not relocate the advanced 3-D object recognition or interpretation to V1. This conjecture is not, therefore, in conflict with the research that locates object 3-D representation further along the processing hierarchy.

Perhaps more philosophically, this conjecture dispenses with the concept of quantity encoding by neurons to be used somehow at later stages of processing. The latter is a mathematical/algorithmic concept, not a circuit concept. It requires a degree of precision and range in numerical representation which seems impractical for neurons, and even more taxing, the quantitative computational mechanisms necessary to make use of such number-like representations. I assume this category of objection has been raised before, but if it has, it is most often ignored in interpreting experimental results. By contrast in the conjecture presented here, disparity is never captured as a quantity in neurons, and does not have to be combined mathematically with a quantified capture of vergence angle to compute distance. The shift required to make a match is selected by pattern matching, but its magnitude is never stored. The shift is used directly to build the 3-D reconstruction, which is disparity’s only use. Implied here is that the 3-D reconstruction is passed along to higher stages for further processing, with shape and location already explicitly represented.

